# Behavioral correlates of cortical semantic representations modeled by word vectors

**DOI:** 10.1101/2020.08.06.240705

**Authors:** Satoshi Nishida, Antione Blanc, Naoya Maeda, Masataka Kado, Shinji Nishimoto

## Abstract

The quantitative modeling of semantic representations in the brain plays a key role in understanding the neural basis of semantic processing. Previous studies have demonstrated that word vectors, which were originally developed for use in the field of natural language processing, provide a powerful tool for such quantitative modeling. However, whether semantic representations revealed by the word vector-based models actually capture our perception of semantic information remains unclear, as there has been no study explicitly examining the behavioral correlates of the modeled semantic representations. To address this issue, we compared the semantic structure of nouns and adjectives estimated from word vector-based brain models with that evaluated from human behavior. The brain models were constructed using voxelwise modeling to predict the functional magnetic resonance imaging (fMRI) response to natural movies from semantic contents in each movie scene through a word vector space. The semantic dissimilarity of word representations was then evaluated using the brain models. Meanwhile, data on human behavior reflecting the perception of semantic dissimilarity between words were collected in psychological experiments. We found a significant correlation between brain model- and behavior-derived semantic dissimilarities of words. This finding suggests that semantic representations in the brain modeled via word vectors appropriately capture our perception of word meanings.

**Author summery:** Word vectors, which have been originally developed in the field of engineering (natural language processing), have been extensively leveraged in neuroscience studies to model semantic representations in the human brain. These studies have attempted to model brain semantic representations by associating them with the meanings of thousands of words via a word vector space. However, there has been no study explicitly examining whether the modeled semantic representations actually capture our perception of semantic information. To address this issue, we compared the semantic representational structure of words estimated from word vector-based brain models with that evaluated from behavioral data in psychological experiments. The results revealed a significant correlation between these model- and behavior-derived semantic representational structures of words. This indicates that the brain semantic representations modeled using word vectors actually reflect the human perception of word meanings. Our findings contribute to the establishment of word vector-based brain modeling as a useful tool in studying human semantic processing.

## 1. Introduction

Natural language processing is a branch of machine learning that aims to develop machines that understand the meanings of words. In the field of natural language processing, a number of algorithms have been developed to capture the semantic representations of words from word statistics in large-scale text data as word vectors [1–5]. The word vectors obtained using these algorithms have effectively captured the latent semantic structure of words and further performed various types of natural language tasks, such as word similarity judgment [3,4,6], sentiment analysis [7,8], and question answering [9].

Furthermore, word vectors can be also used in neuroimaging studies to model semantic representations in the brain [10–17]. These studies have reported that word vector-based models have the ability to predict the brain response evoked by semantic perceptual experiences [10,12–16]. These models are also able to recover perceived semantic contents from brain response [11,17,18]. These findings suggest that word vectors capture at least some aspects of the semantic representations in the brain. However, whether the semantic representations modeled by word vectors accurately reflect the semantic perception of humans is yet to be determined. In other words, no study has yet identified the behavioral correlates of the modeled semantic representations. This clarification is important in order to establish the brain modeling with word vectors as an accurate methodology for investigating human semantic processing.

To examine the behavioral correlates of the modeled semantic representations, we compared the semantic representational structure estimated by word vector-based models with that evaluated from human behavior. The modeled semantic representational structure was obtained using voxelwise modeling with a word vector space of fastText [4] (Fig 1a). This voxelwise model predicts functional magnetic resonance imaging (fMRI) signals evoked by natural movies from the semantic contents in individual movie scenes in the fastText vector space. We then transformed fastText word vectors into brain representations using the model weights, and a brain-derived word dissimilarity matrix from these brain representations was obtained (Fig 1b). Meanwhile, the behavior-derived semantic representational structure was obtained from behavioral data in a psychological task, in which participants separately arranged tens of words (nouns and adjectives) in a two-dimensional space as per their semantic relationship (Fig 2). This task was a modified version of a psychological task introduced previously [19]. A behavior-derived word dissimilarity matrix was then estimated using these behavioral data. Finally, we examined the correlation between the brain- and behavior-derived word dissimilarity matrices separately for both nouns and adjectives.

**Fig 1.**
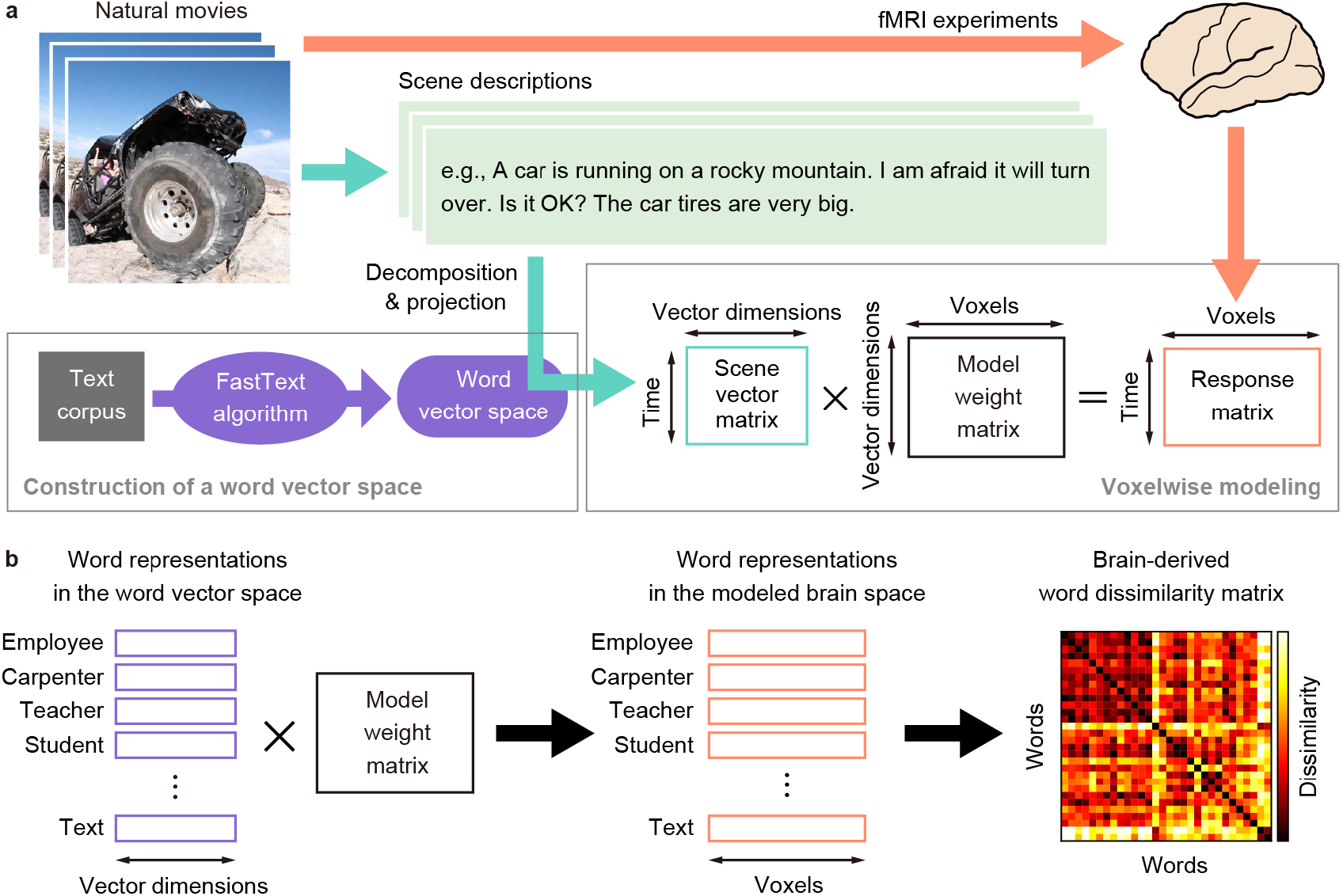
Voxelwise modeling and brain-derived word dissimilarity. **a**) Voxelwise modeling based on a fastText vector space. The model predicts fMRI signals evoked by natural movie scenes via a weighted linear summation of the vector representations of semantic descriptions of each scene (scene vectors). Scene vectors were obtained by transforming manual descriptions of each movie scene through a fastText vector space pretrained using a text corpus. The weights of the linear prediction model were trained using the corresponding time series of movies and fMRI signals of each brain. **b**) Estimation of brain-derived word dissimilarity. Word representations in the fastText vector space were transformed into word representations in the modeled brain space by multiplying fastText word vectors by the model weights. Then, the correlation distance of the modeled word representations between all possible pairs of words was calculated, producing a brain-derived word dissimilarity matrix.

**Fig 2.**
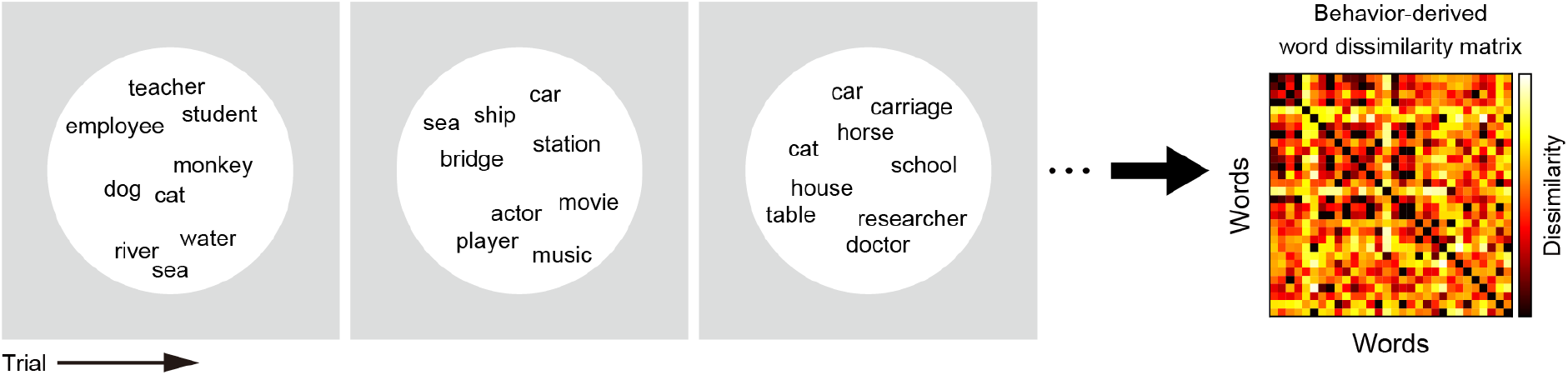
Word-arrangement task and behavior-derived word dissimilarity. To evaluate the word dissimilarity structure derived from human behavior, we conducted psychological experiments in which each participant performed a word-arrangement task. On each trial of this task, participants were required to arrange multiple (≤60) words in a two-dimensional space according to the semantic relationship of those words. After each participant completed ≤1 h of this task, a behavior-derived word dissimilarity matrix was established using inverse multidimensional scaling (see Methods for more details).

## 2. Results

### 2.1. Performance of voxelwise models based on word vectors

We first determined whether the voxelwise models based on a fastText vector space appropriately predict movie-evoked brain responses. The performance of brain-response prediction was evaluated using the following two measures: (1) prediction accuracy calculated as the correlation coefficients between predicted and measured brain responses in the test dataset, and (2) the fraction of significant voxels that reach their prediction accuracy to a threshold value (p < 0.05 after the correction for multiple comparisons using the false discovery rate [FDR]). The prediction accuracy averaged across all cortical voxels and across participants was 0.0928 for movie set 1 (28 participants) and 0.0865 for movie set 2 (40 participants); no participant showed mean prediction accuracy less than 0 (Fig 3a and c). This accuracy was sufficiently high because it was averaged over all cortical voxels, and previous studies on voxelwise modeling have reported a similar tendency [20–22]. The fraction of significant voxels averaged across participants was 0.575 for movie set 1 and 0.526 for movie set 2 (Fig 3b and d). These results indicate that the voxelwise models trained showed sufficient performance in the modeling of semantic representations consistently across different stimulus sets.

**Fig 3.**
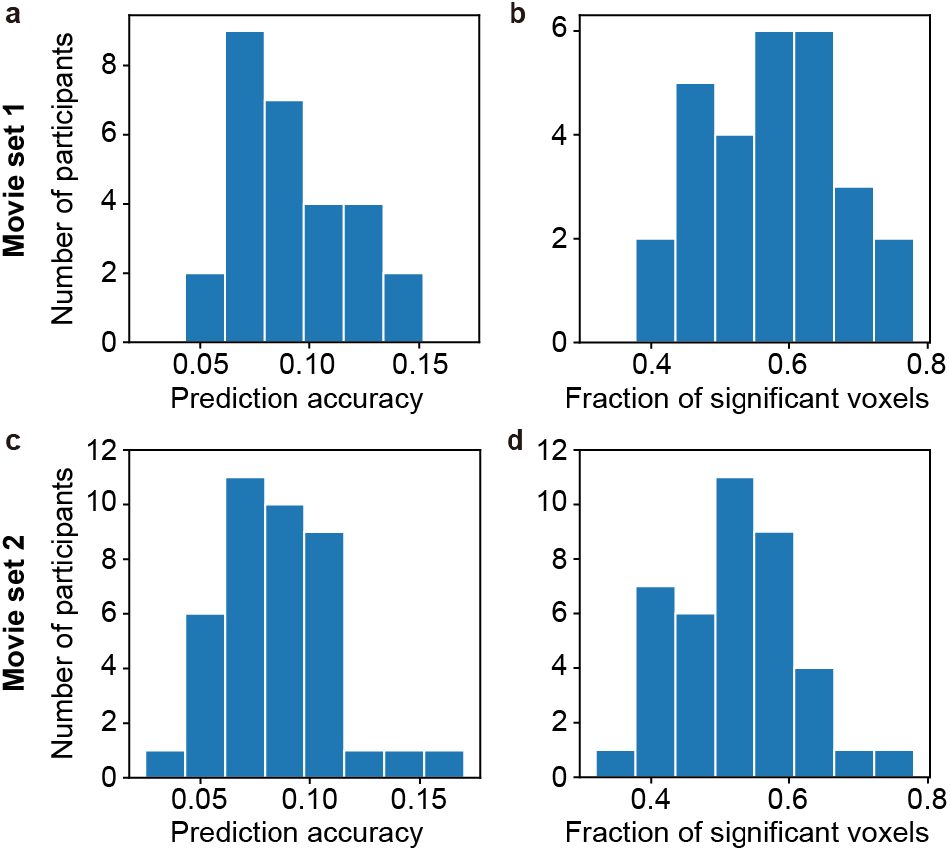
Performance of voxelwise models in brain-response prediction. The performance of voxelwise models (vector dimension = 1000) was evaluated in terms of predicting brain responses in the test dataset. For this purpose, prediction accuracy and the fraction of significant voxels were calculated for each brain. The distribution of prediction accuracy (**a** and **c**) and that of the fraction of significant voxels (**b** and **d**) were separately shown for movie sets 1 (**a** and **b**) and 2 (**c** and **d**).

Although the vector dimensionality of the fastText vector space had little effect on the model performance, there was no clear tendency toward vector-dimensionality dependency (S1 Fig). The vector space of 100 dimensions has showed the highest performance for movie set 1, whereas the vector space of 2000 dimensions showed the highest performance for movie set 2. In addition, the change of model performance was not monotonic against the change of vector dimensions.

### 2.2. Localization of cortical regions highly predicted by voxelwise models

We next identified which cortical regions were predictable by the word vector-based models in order to determine that the models could capture the semantic representations in appropriate cortical regions. For this purpose, the prediction accuracy averaged across participants was calculated in each of anatomically segmented cortical regions and was mapped onto the cortical surface of a reference brain (Fig 4a–b). The highly predictable regions were localized over widespread cortical regions, including the occipital, superior and inferior temporal, and posterior parietal regions, for both movie sets. This localization was consistent with previous reports, in which the semantic representations in the brain were modeled with word features [12,23]. There was a strong correlation of mean prediction accuracy over 150 cortical regions between movie sets 1 and 2 (Pearson’s r = 0.974); this indicates the consistency of model predictability of these cortical regions across stimulus sets. In addition, there were similar tendencies across vector dimensionality (S2 Fig and S1 Table). These results indicate that the voxelwise models trained reliably showed the localization of highly predictable regions.

**Fig 4.**
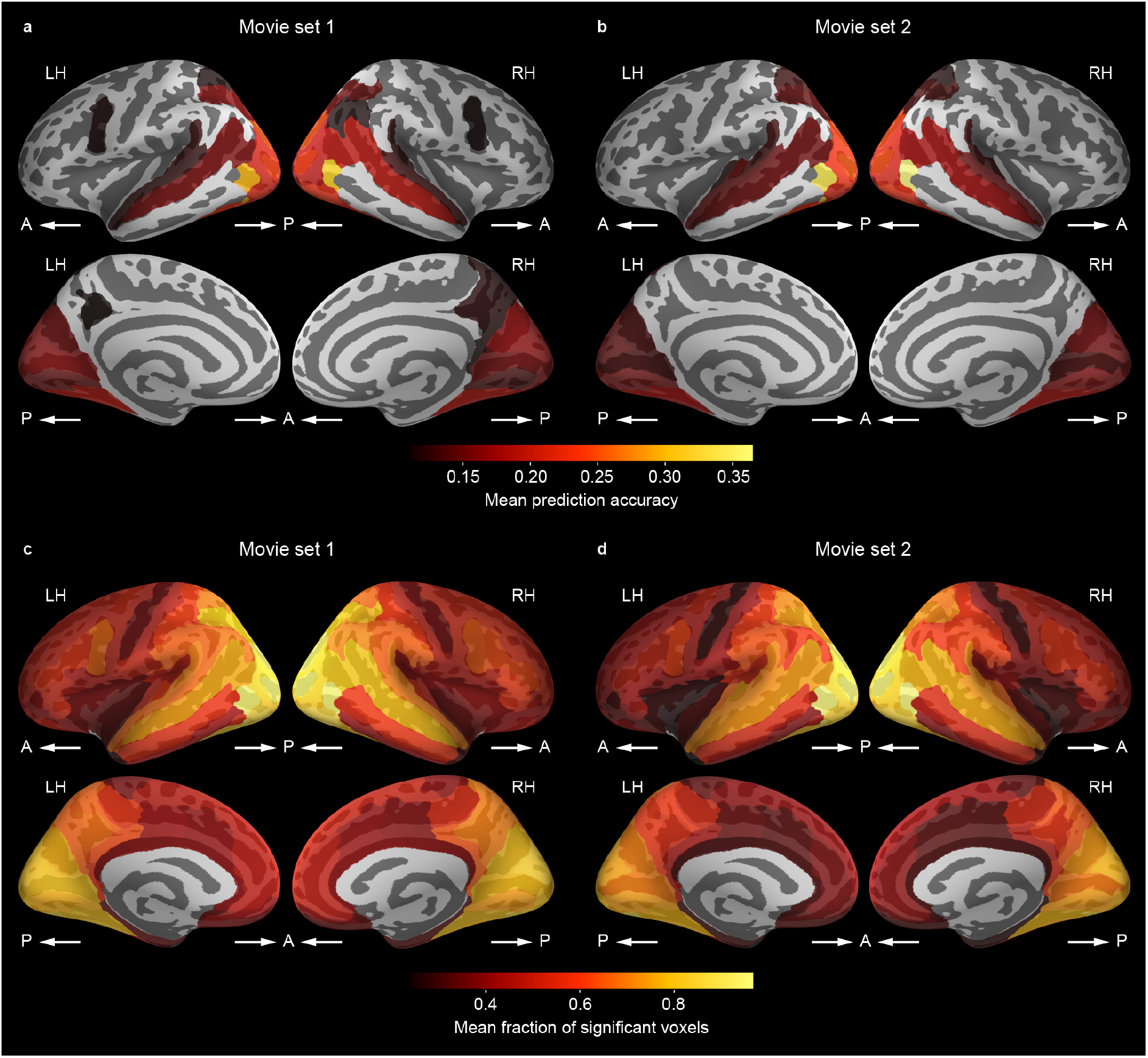
Cortical mapping of model performance. **a**,**b**) Participant-averaged prediction accuracy of voxelwise models (vector dimension = 1000) mapped onto the cortical surface of a reference brain for each of the movie sets 1 (**a**) and 2 (**b**). The prediction accuracy was averaged within each of the cortical regions that were anatomically segmented. Brighter colors in the surface maps indicate cortical regions that have higher prediction accuracy. Only regions with mean prediction accuracy above 0.11 (corresponding to p = 0.0001 ∼ 0.05/150 regions) are shown. **c**, **d**) Fraction of significant voxels of the same models mapped onto the same cortical surface separately for movie sets 1 (**c**) and 2 (**d**). The fraction was then computed within each cortical region. Brighter colors indicate regions that have larger fraction. LH, left hemisphere; RH, right hemisphere; A, anterior; P, posterior.

A similar localization was also observed when the fraction of significant voxels in each cortical region was mapped onto the cortical surface (Fig 4c–d). The fraction was relatively large in the occipital, superior and inferior temporal, and posterior parietal regions compared with the other regions. This localization pattern was observed to be highly consistent across both movie sets and vector dimensions (S3 Fig and S3 Table). Taken together, these results suggest that the word vector-based models capture the semantic representations in appropriate cortical regions.

It should be noted that, however, the fraction of significant voxels was not small even in the other cortical regions, including the prefrontal cortex. The minimum fraction within any individual region was 0.237, which is significantly higher than 0.05 (corresponding to the significance level of 0.05; Wilcoxson test, p < 0.00001). This result suggests that although high prediction accuracy was observed in specific regions (Fig 4a–b and S2 Fig), the word vector-based model can potentially capture semantic information even from other regions across the cortex.

### 2.3. Correlation between brain- and behavior-derived word dissimilarities

Finally, we tested the correlation between the word dissimilarity matrix derived from the voxelwise models and that derived from behavioral data to clarify the behavioral correlates of modeled semantic representations. Word representations in the modeled brain space were calculated by multiplying the fastText word vectors by model weights, and the brain-derived word dissimilarity matrix was obtained from the correlation distance between all possible pairs of word representations (Fig 1b). Meanwhile, the behavior-derived word dissimilarity matrix was measured from the behavioral data from the word-arrangement task (Fig 2), in which 36 participants completed 18.8 (SD = 10.7) trials on average for each session (see Methods for more details).

These brain- and behavior-derived word dissimilarity matrices were constructed by averaging the matrices over all model or behavioral data separately for nouns (Fig 5a) and adjectives (Fig 5b). We found significant correlations between these two types of word dissimilarity matrices. For nouns, the correlation coefficients were consistently high across movie sets (r = 0.612 and 0.635 for movie sets 1 and 2, respectively; p < 0.00001). For adjectives, the correlation coefficients were deemed smaller but significant for both movie sets (r = 0.342 and 0.358 for movie sets 1 and 2; p < 0.00001). In these cases, the populations from which we obtained these dissimilarity matrices were different between brain- and behavior-derived data. However, we observed a significant correlation for both nouns and adjectives and for both movie sets even when these matrices were collected from the same population (p < 0.00001; S4 Fig). In addition, significant correlations were consistently observed regardless of vector dimensionality (S5 Fig a and c); although the correlation coefficients differed across vector dimensions, no clear tendency of changes in correlation coefficients across vector dimensions was determined (S5 Fig b and d). These results indicate that the brain-derived dissimilarity structure of semantic representations correlates with the behavior-derived dissimilarity structure.

**Fig 5.**
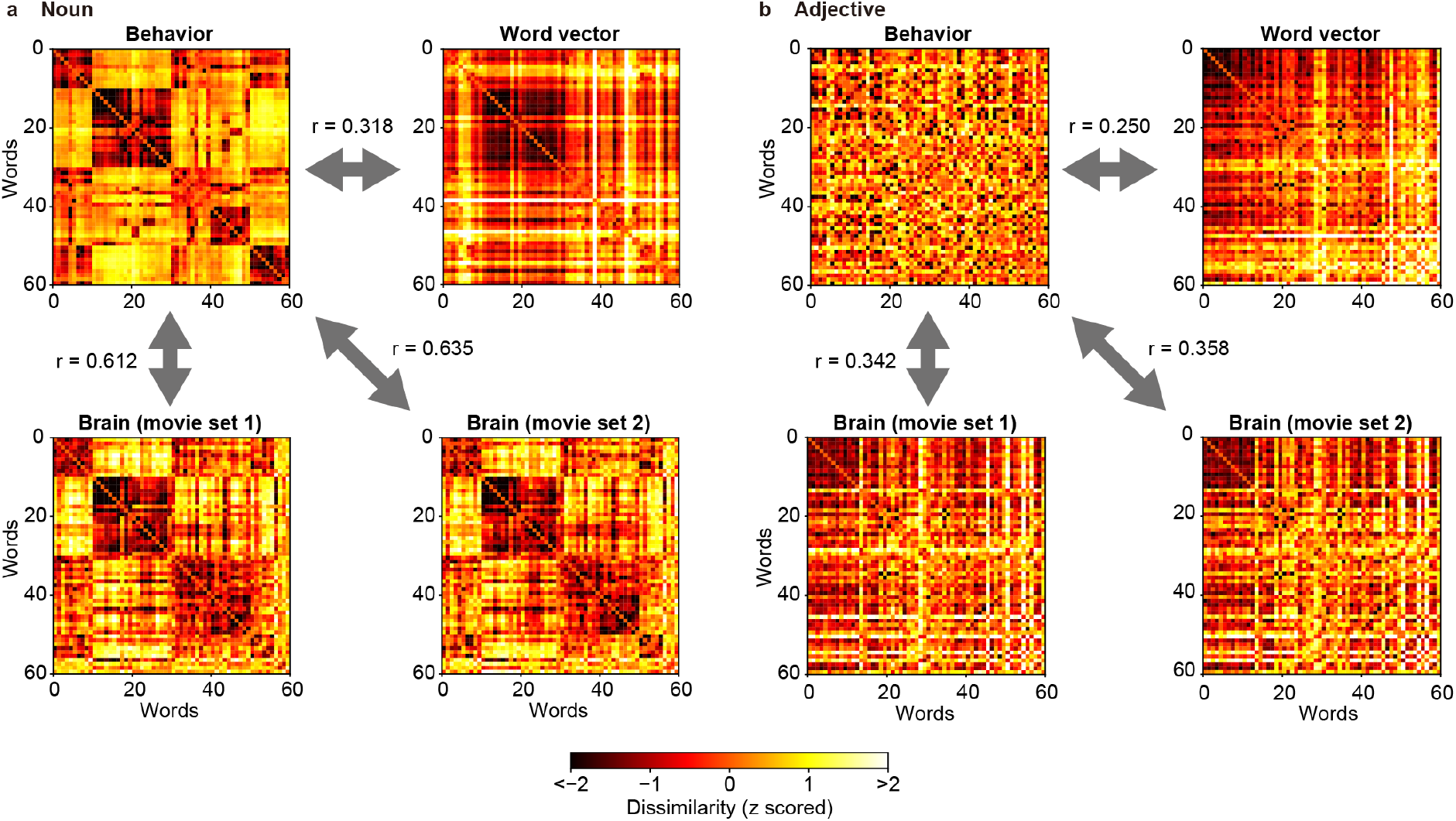
Correlations between dissimilarity matrices. We have constructed word dissimilarity matrices separately for nouns (**a**) and adjectives (**b**). Each color map shows the behavior-derived matrices obtained from behavioral data (top left in **a** and **b**), the word vector-derived matrices obtained directly from the fastText vector space (top right), or the brain-derived matrices obtained from voxelwise models (vector dimension = 1000) for each movie set (bottom). Brighter colors in each map indicate higher dissimilarity of word pairs. Significant correlations were identified between behavior- and word vector-derived matrices and between behavior- and brain-derived matrices (p < 0.0001). Pearson’s correlation coefficients (r) are indicated.

To determine whether the behavioral correlates of word dissimilarity change through the transformation from original fastText word vector representations to brain representations, we evaluated the correlation of a word vector-derived word dissimilarity matrix with the behavior-derived matrix. We found that correlation coefficients between the word vector- and behavior-derived matrices were significantly lower than those between the brain- and behavior-derived matrices (Fig 5; r = 0.318 and 0.250 for nouns and adjectives, respectively; bootstrapping test, p < 0.0001). This observation was consistent across vector dimensions (except for the vector dimension = 100 for adjectives; S5 Fig a and c). These results suggest that the semantic representations of word vectors become more similar to human perception as they are transformed into brain semantic representations.

The brain-derived dissimilarity matrices in Fig 5, S4 and S5 Figs were calculated from the model weights of all cortical voxels (the number of voxels was 54914–73301 [mean ± SD = 62245 ± 5018] for movie set 1 and 52832–73301 [mean ± SD = 62150 ± 5190] for movie set 2). However, the high performance of the voxelwise models was observed in localized cortical regions (Fig 4, S2 and S3 Figs). Hence, the correlation of brain- and behavior-derived dissimilarity matrices may be stronger when only the cortical regions with high model performance were used. To test this possibility, we constructed brain-derived dissimilarity matrices using only those voxels with the highest prediction accuracy (top 2000, 5000, 10000, 30000, or 50000 voxels). Then, their correlations with behavior-derived matrices were compared with the original correlations calculated using all cortical voxels. However, we found the strongest correlation when using all cortical voxels (Fig 6 and S6 Fig). This result suggests that semantic information correlated with behavior are distributed broadly across the cortex and can be captured using word vector-based voxelwise models.

**Fig 6.**
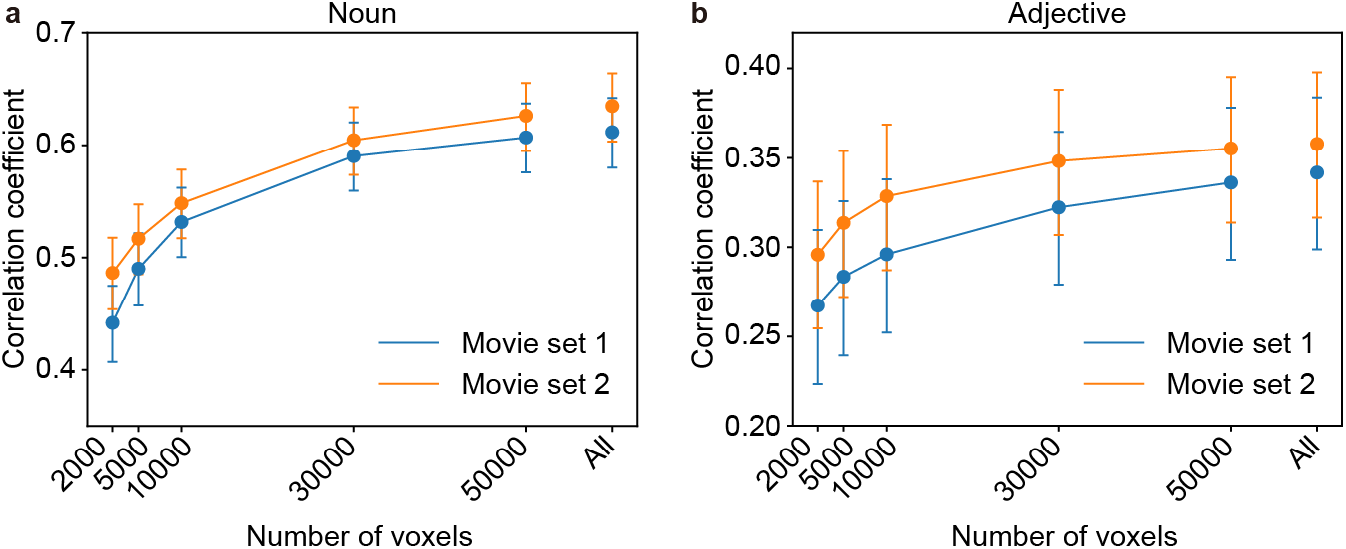
Effects of voxel selection on brain–behavior correlation. We have evaluated the correlations between brain- and behavior-derived word dissimilarity matrices when the brain-derived matrix was obtained from the selected voxels with the highest model accuracy. The correlation coefficients are shown separately for nouns (**a**) and adjectives (**b**), while the numbers of selected voxels are changed (2000, 5000, 10000, 30000, 50000, and all voxels). Error bars indicate 95 % confidence interval (CI) estimated by bootstrapping.

## 3. Discussion

We have examined the behavioral correlates of semantic representations estimated by word vector-based brain models. We constructed a voxelwise model that has the ability to predict movie-evoked fMRI signals in individual brains through a fastText vector space. The voxelwise models showed substantial response-prediction performance in reasonable cortical regions consistently across stimulus and parameter sets. There were significant correlations between the word dissimilarity structure derived from the voxelwise models and that derived from behavioral data. These results suggest that the semantic representations estimated from word vector-based brain models can appropriately capture the semantic relationship of words in human perception. Our findings contribute to the establishment of word vector-based brain modeling as a powerful tool in investigating human semantic processing.

Word vectors have been extensively leveraged for the modeling of semantic information in the human brain. Some studies have successfully predicted brain responses, through word vector spaces, evoked by audiovisual stimuli, such as movies [15,16,22], pictures [10,13], and sentences [14]. In addition, one of these studies has visualized the semantic representational space in the brain through word vector-based modeling [16]. Another line of studies has successfully recovered perceived semantic contents from brain responses by using word vector spaces [11,17,18]. In this way, word vectors will be a useful tool to investigate comprehensively the semantic processing in the human brain. However, no study has explicitly clarified the behavioral correlates of the brain semantic representations revealed by word vector-based models. Although previous studies in which movie-evoked brain responses were predicted from manual descriptions for the movie scenes [11,15] employed human behavior implicitly, they did not reveal the association between brain- and behavior-derived semantic representations. To the best of our knowledge, this is the first study demonstrating the behavioral correlates of semantic representations modeled by word vectors.

One of the advantages of word vector-based modeling is that semantic representations can be quantified using much more vocabularies [11] than modeling based on discrete word features (e.g., [23]). This allows the quantification of different aspects of semantic information in the brain by using different types of words. In particular, adjectives have a potential for the visualization of impression information [11]. We found the behavioral correlates of modeled semantic representations on nouns and even adjectives, which were consistently observed across stimulus and parameter sets (Fig 5, S4 and S5 Figs). These findings indicate that word vectors are deemed suitable for modeling the cortical representations of impression information. Thus, word vector-based modeling provides a useful framework for extensive investigation of semantic representations in the brain.

In this study, we found localized cortical regions in which brain responses were accurately predicted by our models (Figs 2 and 4, S2 and S3 Figs). Although these brain regions, primarily the posterior and temporal cortices, are consistent with previous reports on the modeling of semantic representations [12,23], it may be argued that these regions do not encompass the entire language network, including the left inferior frontal and superior temporal cortices [24]. However, a recent study has clearly demonstrated that language network is not necessarily involved in semantic processing that requires no explicit linguistic processing [25]. They reported that aphasia patients with physical damage to the language network manifested severely declined performance in semantic tasks with linguistic information, but performance equivalent to healthy controls in semantic tasks with nonlinguistic information. This notion is also supported by evidence that semantic models using word features can predict brain response in the language network when the response is evoked by linguistic stimuli, e.g., sentences and speech [14,21,26]; however, it was not applicable when the response is evoked by nonlinguistic stimuli, e.g., movies [12,23]. Thus, the semantic representations modeled using the brain response evoked by nonlinguistic stimuli may be biased toward specific modalities, such as vision and audition.

In the field of natural language processing, recently developed algorithms based on deep learning have exhibited state-of-the-art performance in many types of natural language tasks [27–29]. The feature representations obtained by these algorithms are also beginning to be used for the modeling of semantic representations [30,31], which show state-of-the-art performance even in brain-response prediction [32]. Hence, it is possible that the correlation between brain- and behavior-derived semantic representations improves by using such deep learning features instead of word vectors. However, some advantages are noted using word vectors compared with deep learning features: First, word vectors are easily trained with much smaller sets of parameters than deep learning features. Second, the internal processing of word vector algorithms is more explainable than that of deep learning-based algorithms. Third, there are many existing methods to improve word-vector spaces for specific types of tasks [13,33–37]. These advantages may be also beneficial for the modeling of semantic representations. Therefore, it is important to use these two types of algorithms separately depending on the aims of specific studies on semantic processing.

## 4. Methods

### 4.1. Participants

In total, 52 healthy Japanese participants (21 females and 31 males; age 20–61, mean ± SD = 26.8 ± 8.9 years) were recruited for the 2 sets of fMRI experiments. Among the recruits, 36 participated in one or the other of these fMRI experiments, and 16 participated in both. In addition, another 36 healthy Japanese participants (20 females and 16 males; age 18–58, mean ± SD = 25.9 ± 10.1 years) were recruited for psychological experiments. Of these, 6 were also participants of the fMRI experiments and 30 were unique participants. All participants had normal or corrected-to-normal vision. Written informed consent was obtained from all of the participants. The experimental protocol was approved by the ethics and safety committees of the National Institute of Information and Communications Technology. The fMRI data, but not the psychological data, used here were also utilized in a previous publication [22].

### 4.2. MRI experiments

Functional and anatomical MRI data were collected via a 3T Siemens MAGNETOM Prisma scanner (Siemens, Germany), with a 64-channel Siemens volume coil. Functional data were collected using a multiband gradient echo EPI sequence [38] (TR = 1000 ms; TE = 30 ms; flip angle = 60 °; voxel size = 2 × 2 × 2 mm; matrix size = 96 × 96; FOV = 192 × 192 mm; the number of slices = 72; multiband factor = 6). Anatomical data were also gathered using a T1-weighted MPRAGE sequence (TR = 2530 ms; TE = 3.26 ms; flip angle = 9 °; voxel size = 1 × 1 × 1 mm; matrix size = 256 × 256; FOV = 256 × 256 mm; the number of slices = 208) on the same 3T scanner.

In the two sets of fMRI experiments, participants were asked to view movie stimuli on a projector screen inside the scanner (27.9 × 15.5 of visual angle at 30Hz) and used MR-compatible headphones for the sounds. The participants were given no explicit task. The fMRI data for each participant upon viewing of the movies were collected in three separate recording sessions over 3 days for each set of fMRI experiments.

The movie stimuli consisted of Japanese television advertisements for one set of experiments (movie set 1) and Japanese web advertisements for the other set of experiments (movie set 2; see also [22]). Movie set 1 has included 420 ads broadcasted on Japanese TV between 2011 and 2017; meanwhile, movie set 2 included 368 ads broadcasted on the Internet between 2015 and 2018. The ad movies were all unique, include a wide variety of product categories (see S3 Table for more details). The length of each movie was 15 or 30 s. To create the movie stimuli for each experiment, the original movies in each of the movie sets 1 and 2 were sequentially concatenated in a pseudo-random order. For each movie set, 14 non-overlapping movie clips of 610 s in length were obtained.

Individual movie clips were then displayed in separate scans. The initial 10 s of each clip served as a dummy in order to discard hemodynamic transient signals caused by clip onset. fMRI responses collected during the 10 s dummy part were not used for modeling. Twelve clips from each movie set were only presented once. The fMRI responses to these clips were used for the training of the voxelwise models (training dataset; 7200 s in total). The other two clips for each movie set were presented four times each in four separate scans. The fMRI responses to these clips were then averaged across four scans, which were further used for the test of the voxelwise models (test dataset; 1200 s in total).

### 4.3. MRI data pre-processing

Motion correction in each functional scan was carried out via the statistical parameter mapping toolbox (SPM8, http://www.fil.ion.ucl.ac.uk/spm/software/spm8/). For each subject, all volumes were aligned to the first image from the first functional run. Low-frequency fMRI response drift was then eliminated by subtracting median-filtered signals (within a 120-s window) from raw signals. Then, the response for each voxel was normalized by subtracting the mean response and scaling to the unit variance. FreeSurfer [39,40] was used to identify cortical surfaces from anatomical data and register these surfaces to the voxels of functional data. Then, each voxel was assigned to one cortical region based on the atlas for cortical segmentation [41].

### 4.4. Word vector spaces

This study used the skip-gram algorithm of fastText [4] to construct a word vector space. This algorithm has been originally developed to learn a word vector space based on nearby word co-occurrence statistics in natural language texts. The training objective of the skipgram algorithm is to obtain word vector representations that enable the surrounding words to be accurately predicted from a given word in a sentence. More formally, given a sequence of training words *w*_1_, *w*_2_,…, *w_T_*, the skip-gram algorithm seeks a *K*-dimensional vector space that maximizes the average log probability, given as:

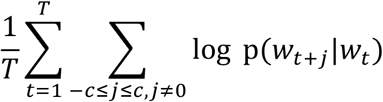

where *c* is the size of the context window, which corresponds to the number of to-be-predicted words before and after the center word *w_t_*. Therefore, the skip-gram vector space is optimized on the basis of the local co-occurrence statistics of nearby words in the text corpus. The basic formulation of *p*(*w_t+j_*|*w_t_*) is the softmax function:

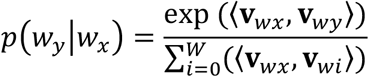

where ***v***_*wi*_ is the vector representation of *w_i_*, *W* is the number of words in the vocabulary, and 〈**v**_1_,**v**_2_〉 indicates the inner product of vectors **v**_1_ and **v**_2_. However, because of the high computational cost of this formulation, the negative sampling technique is used to produce a computationally efficient approximation of the softmax function [2]. In addition, fastText introduces sub-word modeling, which is robust for inflected and rare words [4].

A fastText vector space was constructed from a text corpus of the Japanese Wikipedia dump on April 1, 2020. All Japanese texts in the corpus were segmented into words and lemmatized using MeCab (http://taku910.github.io/mecab). Used were only nouns, verbs, and adjectives. In an attempt to improve the reliability of fastText learning, the vocabulary size was restricted to ∼100,000 words by excluding words that appeared fewer than 262 times in the corpus. The learning parameters of fastText used were as follows: window size = 10; the number of negative samples = 5; downsampling rate = 0.001; and the number of learning epochs = 10. The fastText vector spaces were constructed using five different numbers of vector dimensions (100, 300, 500, 1000, and 2000). Of these, the 1000-dimension vector space was used for the main analysis (Figs 3–6), and the other dimensions were used to test the effect of vector dimensionality on modeling performance and behavioral correlates. To eliminate the effects of trial variations in the learning quality of word vector spaces on the performance of voxelwise modeling, five different fastText spaces were learned independently using the same text corpus and the same parameters. All results were obtained from the average over five learned spaces for each parameter set.

### 4.5. Movie scene descriptions

Manual scene descriptions using natural Japanese language were provided for every 1-s scene of each movie in movie sets 1 and 2, in a manner similar to that described previously [11,12,22]. The annotators were native Japanese speakers (movie set 1: 68 females and 28 males, age 19–62 years; movie set 2: 11 females and 2 males, age 20–56 years), who were not the fMRI participants. They were instructed to describe each scene (the middle frame of each 1-s clip) using more than 50 Japanese characters. Multiple annotators (movie set 1, 12–14 annotators; movie set 2, 5 annotators) were randomly assigned for each scene to reduce the potential effect of personal bias. The descriptions contain a variety of expressions reflecting not only objective perceptions but also the subjective perceptions of the annotator (e.g., impression, feeling, association with ideas; for more details, see https://osf.io/3hkwd).

Each description for a given scene was also segmented, lemmatized, and decomposed into nouns, verbs, and adjectives via MeCab as described above; then, they were transformed into fastText vectors. The word vectors were then averaged within each description. For each scene, all vectors obtained from the different descriptions were averaged. Through this procedure, a single vector (scene vector) was obtained for each 1-s scene, which was later used for modeling.

### 4.6. Voxelwise modeling

The procedure of voxelwise modeling was similar to that described previously [12]. A series of fMRI responses evoked in individual *N* voxel by a series of *S* movie scenes, represented by a *S* × *N* matrix **R**, was modeled as a weighted linear combination of a *K*-dimensional scene-vector matrix **V** plus isotropic Gaussian noise ***ε***.

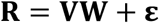

A set of linear temporal filters was used to model the slow hemodynamic response and its coupling with brain response [42]. In an attempt to capture the hemodynamic delay in the responses, the *S* × 4*K* scene-vector matrix was constructed by concatenating four sets of *K*-dimensional scene vectors with delays of 3, 4, 5, and 6 s. The 4*K* × = weighted matrix **W** was estimated using an L2-regularized linear least-squares regression, which can obtain good estimates even for models containing a large number of regressors [23].

In order to estimate the regularization parameters, the training dataset was randomly divided into two subsets containing 80 % and 20 % of the samples, for model fitting and validation, respectively. This random resampling procedure was repeated 10 times. Regularization parameters were optimized, according to the mean Pearson’s correlation coefficient between the predicted and measured fMRI signals for the 20 % validation samples. An optimal parameter was obtained separately for each model.

The brain-response prediction performance of the model was evaluated using the test dataset, which was not used for model fitting or parameter estimation. The prediction accuracy of the model was quantified as the Pearson’s correlation coefficient between the predicted and average measured fMRI signals in the test dataset.

### 4.7. Psychological experiments

Participants were asked to perform a word-arrangement task on a PC. This task is considered to be a modified version of the psychological task introduced previously [19]. Kriegeskorte and colleagues originally used this task to examine the behavioral correlates of cortical object representations, which were quantified using the representational similarity of fMRI signals evoked by object stimuli [43,44]. The present study used this task to study the methodological validity of word vector-based brain modeling by testing the behavioral correlates of modeled cortical semantic representations. The psychological data are available online (https://osf.io/um3qg/).

In each trial of this task, the participants were required to arrange ≤60 words (nouns or adjectives) in a two-dimensional space according to their semantic relationship on a computer screen by mouse drag-and-drop operations (Fig. 2). This paradigm has allowed for the efficient collection of the perceptual semantic dissimilarity between words from participants [19].

The words used in this task included 60 nouns and 60 adjectives in Japanese (Table 1). These words were selected from the vocabulary of the fastText space (i.e., top 100,000 most frequently used words in the Japanese Wikipedia corpus). Nouns were selected in terms of the following six semantic categories: humans, non-human animals, non-animal natural things, constructs, vehicles, and other artifacts. Ten words were selected for each category. For adjectives, such category-based selection was deemed difficult due to the small number of adjectives in the vocabulary (only 473 words). Nouns and adjectives were separately used in two distinct sessions of the task, which was performed over 2 days.

**Table 1.**
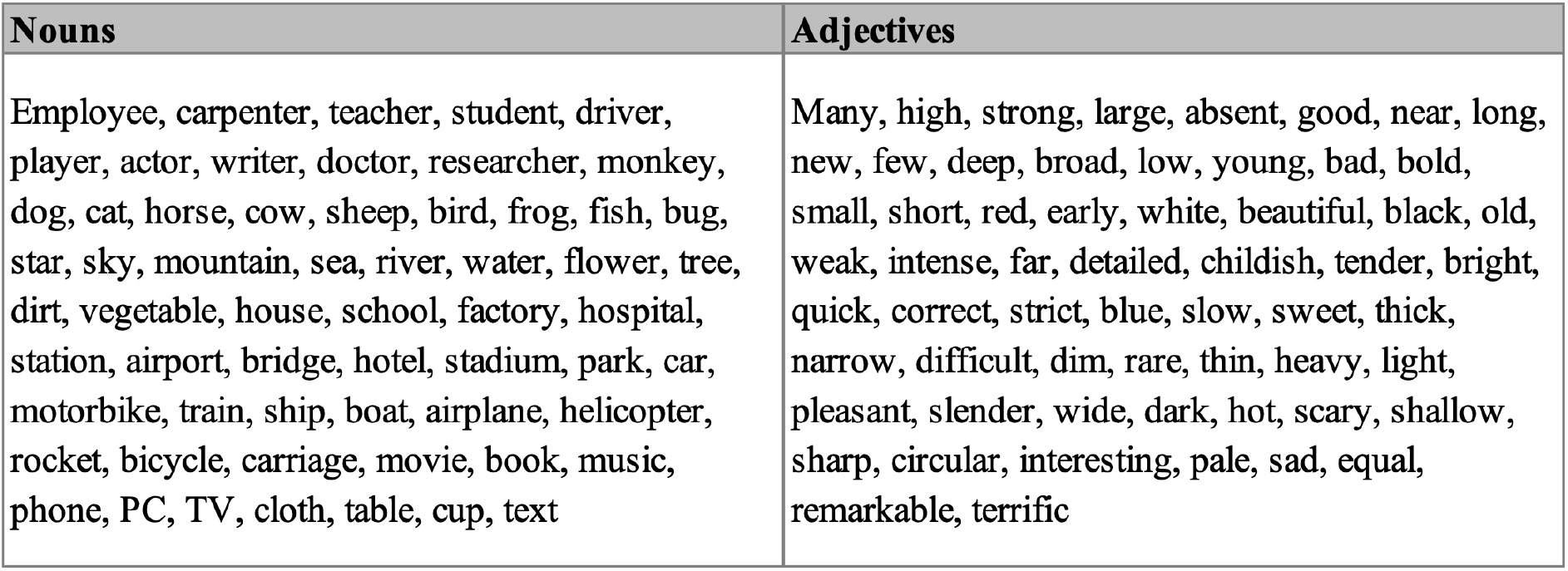
Words used in psychological experiments and for calculating word dissimilarity matrices. Original words were in Japanese, and, for display purposes, here they were translated into English.

The words were arranged in a designated circular area on the computer screen (“arena”). The words were initially displayed outside the arena. The participants used mouse drag-and-drop operations in moving each word item into the arena and arranging them. The arrangement of words was according to the participants’ own judgment of semantic similarity and dissimilarity between words. More similar or dissimilar word pairs should be closer or further apart in the arena, respectively. The participants were allowed to move any words within the arena as many times as they wished. Once all words were moved into the arena, the participants could click a button marked “Next” anytime to move on to the next trial.

On the initial trial of each session, the participants have arranged the entire set of 60 words (nouns or adjectives). On subsequent trials, they arranged subsets of those 60 words. After the end of each trial, the rough estimate of the word dissimilarity matrix and the evidence (0–1) for each word pairwise dissimilarity were computed as described previously [19]. Words in the subsets chosen for each trial were determined so as to increase evidence for pairwise dissimilarities of the words whose evidence estimated on the previous trial was the weakest. The session continued until the evidence of every word pair dissimilarity was above a threshold (0.75) or until the total duration of the session approached 1 h.

### 4.8. Word dissimilarity matrix

The brain-derived word dissimilarity matrix was estimated using the following procedure: First, the 4*K* × = voxelwise model weights for each participant were transformed into *K* × *N* by averaging the weights across the four sets of hemodynamic delay terms. Second, all *N* or top *M* voxels with the highest prediction accuracy were then selected from the model weights. All voxels were used in the main analysis, and selected voxels (*M* = 2000, 5000, 10000, 30000, and 50000) were used in an additional analysis that tested the effect of voxel selection on brain-behavior correlation. Third, the *K*-dimensional fastText vectors for the words used in the psychological experiments (Table 1) were multiplied by the *K* × *N* weight matrix, which yielded *N*- or *M*-dimensional word representations in the modeled brain space. Finally, a word dissimilarity matrix was calculated by the correlation distance (1 –Pearson’s correlation coefficient) between these word representations, which was averaged across participants separately for nouns and adjectives.

The behavior-derived word dissimilarity matrix was evaluated from multiple word-subset arrangements on the word-arrangement task. The dissimilarity for a given word pair was estimated as a weighted average of the scale-adjusted dissimilarity estimates from individual arrangements as has been described previously [19]. These estimates produced a word dissimilarity matrix for the entire set of words. In this way, the word dissimilarity matrix was estimated separately for nouns and adjectives and was averaged across participants.

In addition, a word vector-derived word dissimilarity matrix was also examined in order to test whether the behavioral correlates of semantic dissimilarity structures changed through the transformation from original word vector representations to brain word representations. The word vector-derived matrix was computed by the correlation distance between fastText vectors of the same word sets (Table 1) separately for nouns and adjectives.

The behavioral correlates of brain- or word vector-derived data were evaluated using Pearson’s correlation of the brain- or word vector-derived word dissimilarity matrix with the behavior-derived matrix. The lower triangular portion of each matrix was used for calculating correlation coefficients. A bootstrapping test was performed to statistically compare correlation coefficients. In each repetition of this test, word pairs were randomly resampled from each word dissimilarity matrix with replacement. Then, the difference of correlation coefficients between different pairs of word dissimilarity matrices was calculated. This procedure was repeated 10,000 times to obtain the distribution of correlation coefficient differences between different matrix pairs and estimate the p values.

## Acknowledgments

The work was supported by JSPS KAKENHI Grant-in-Aid for Early-Career Scientists (18K18141) and for Young Scientists A (15H05311), and JST ERATO (JPMJER1801) from MEXT. We thank Ms. Hitomi Koyama, Mr. Yusuke Nakano, Mr. Koji Takashima, Mr. Takeshi Matsuda, Mr. Susumu Minamiyama, Ms. Mami Yamashita, Mr. Ryo Yano, Ms. Risa Matsumoto, Mr. Masato Okino, and Mr. Akira Nagaoka for their analytical and experimental support.

---

**S1 Fig.**
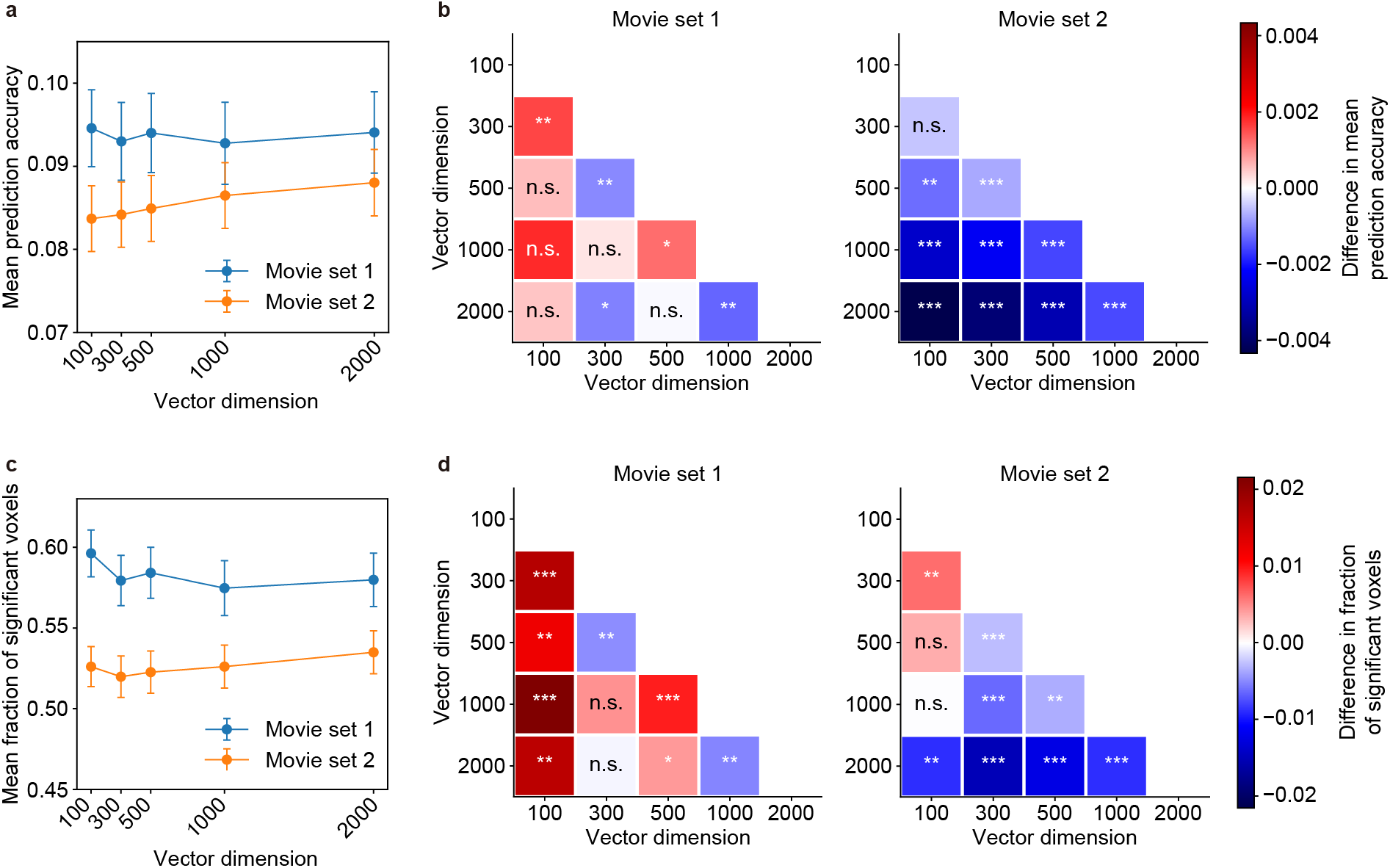
Model performance comparison across vector dimensions. **a**) Mean prediction accuracy of voxelwise models with different vector dimensions. Error bars indicate standard error of the mean (SEM). **b**) Difference of mean prediction accuracies between different vector dimensions. The difference was evaluated separately for movie sets 1 (left) and 2 (right). The color of each cell represents the accuracy difference of the dimension on the x-axis minus the dimension on y-axis (red, positive values; blue, negative values). The mark in each cell indicates the statistical significance of the difference (Wilcoxon test, ***p < 0.0001, **p < 0.01, *p < 0.05, n.s., p > 0.05, FDR corrected). **c**) Fractions of significant voxels for voxelwise models with different vector dimensions. The same conventions were used as in **a**. **d**) Difference in fractions of significant voxels between different vector dimensions. The same conventions were used as in **b**.

**S2 Fig.**
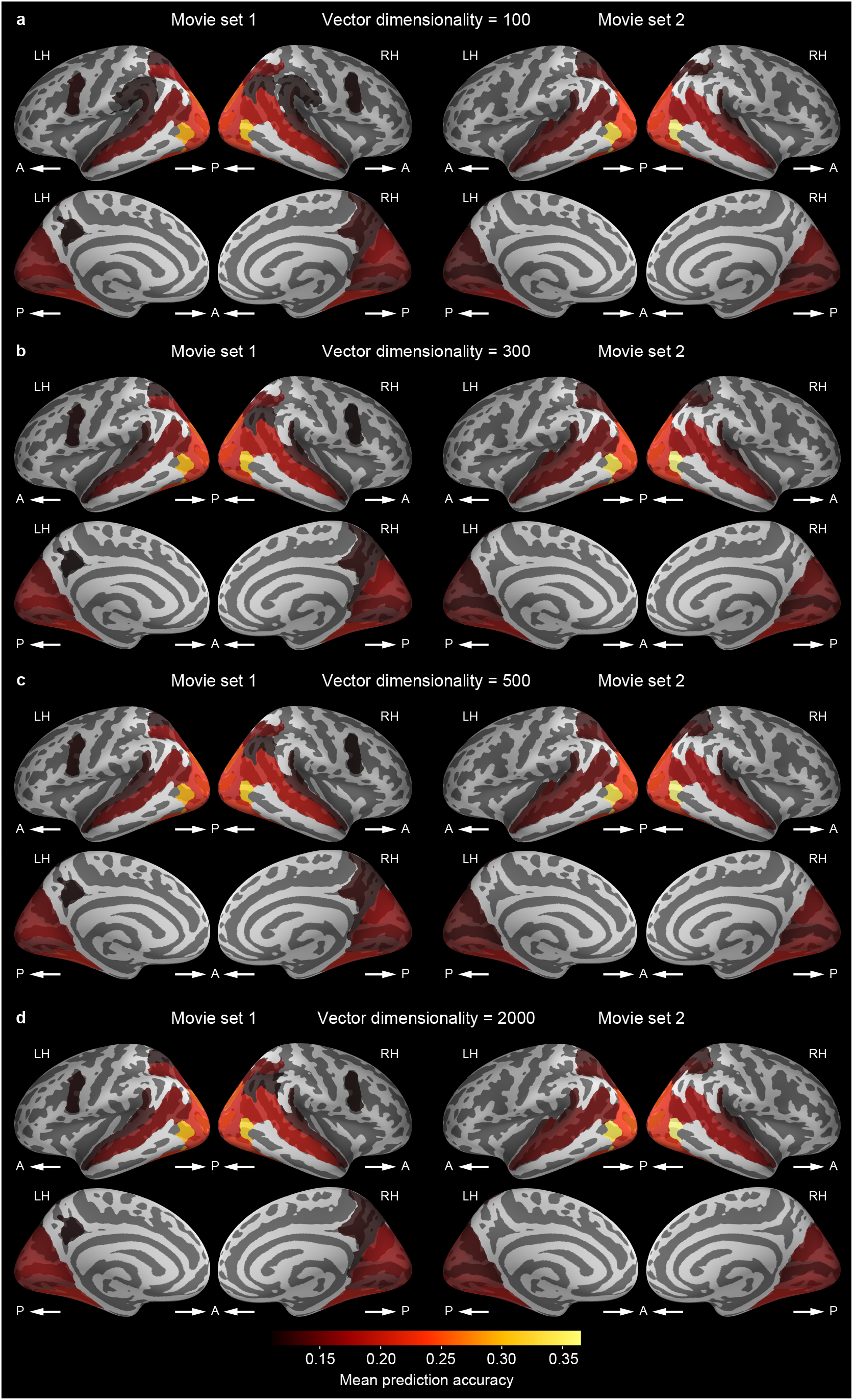
Cortical mapping of prediction accuracy for other vector dimensions. The mean prediction accuracy of voxelwise models in each brain region was mapped onto the cortical surface (**a**, vector dimension = 100; **b**, 300; **c**, 500; **d**, 2000). The same conventions were used as in Fig 4**a**–**b**.

**S3 Fig.**
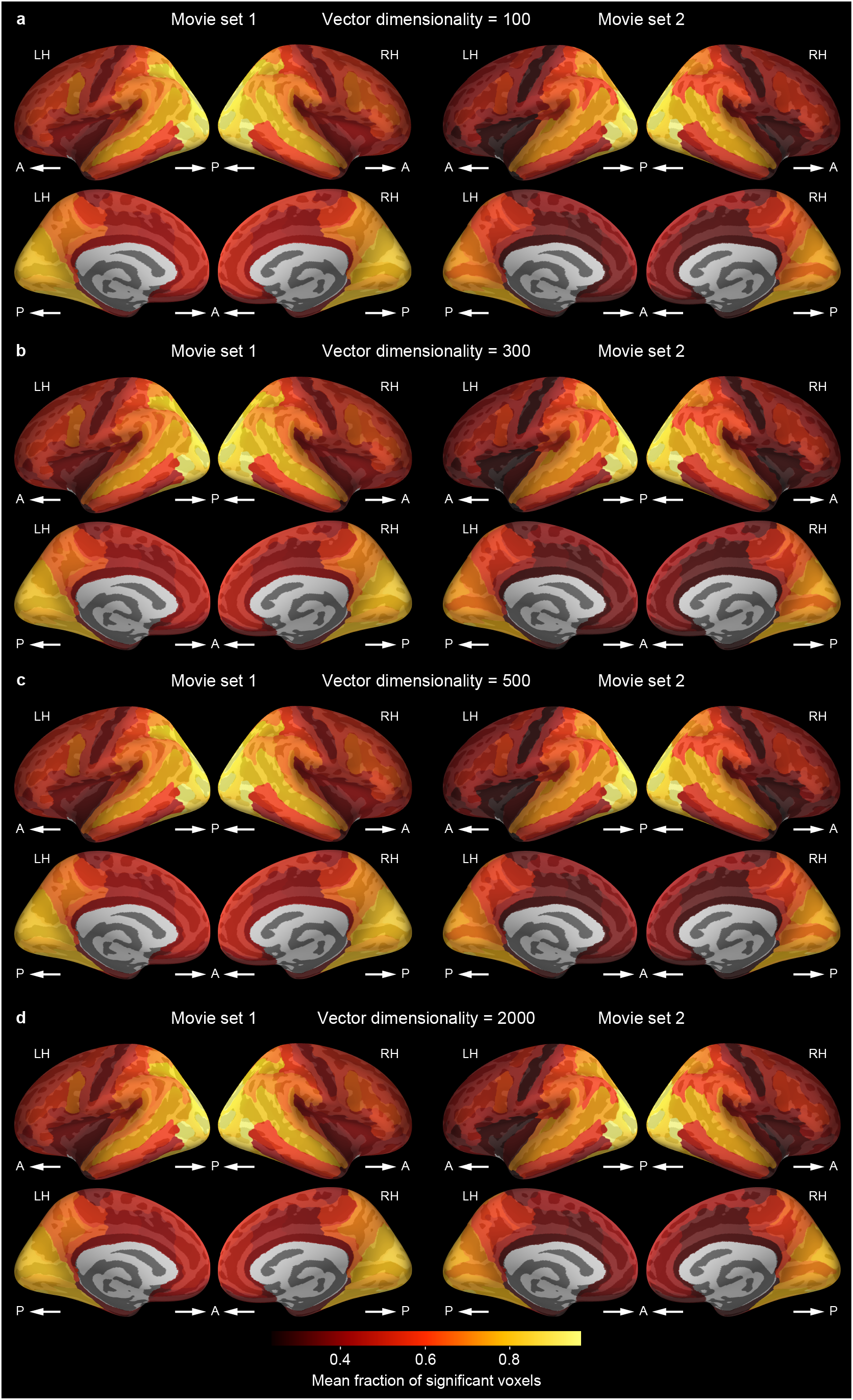
Cortical mapping of the fraction of significant voxels for other vector dimensions. The fraction of significant voxels in each brain region was mapped onto the cortical surface (**a**, vector dimension = 100; **b**, 300; **c**, 500; **d**, 2000). The same conventions were used as in Fig 4**c**–**d**.

**S4 Fig.**
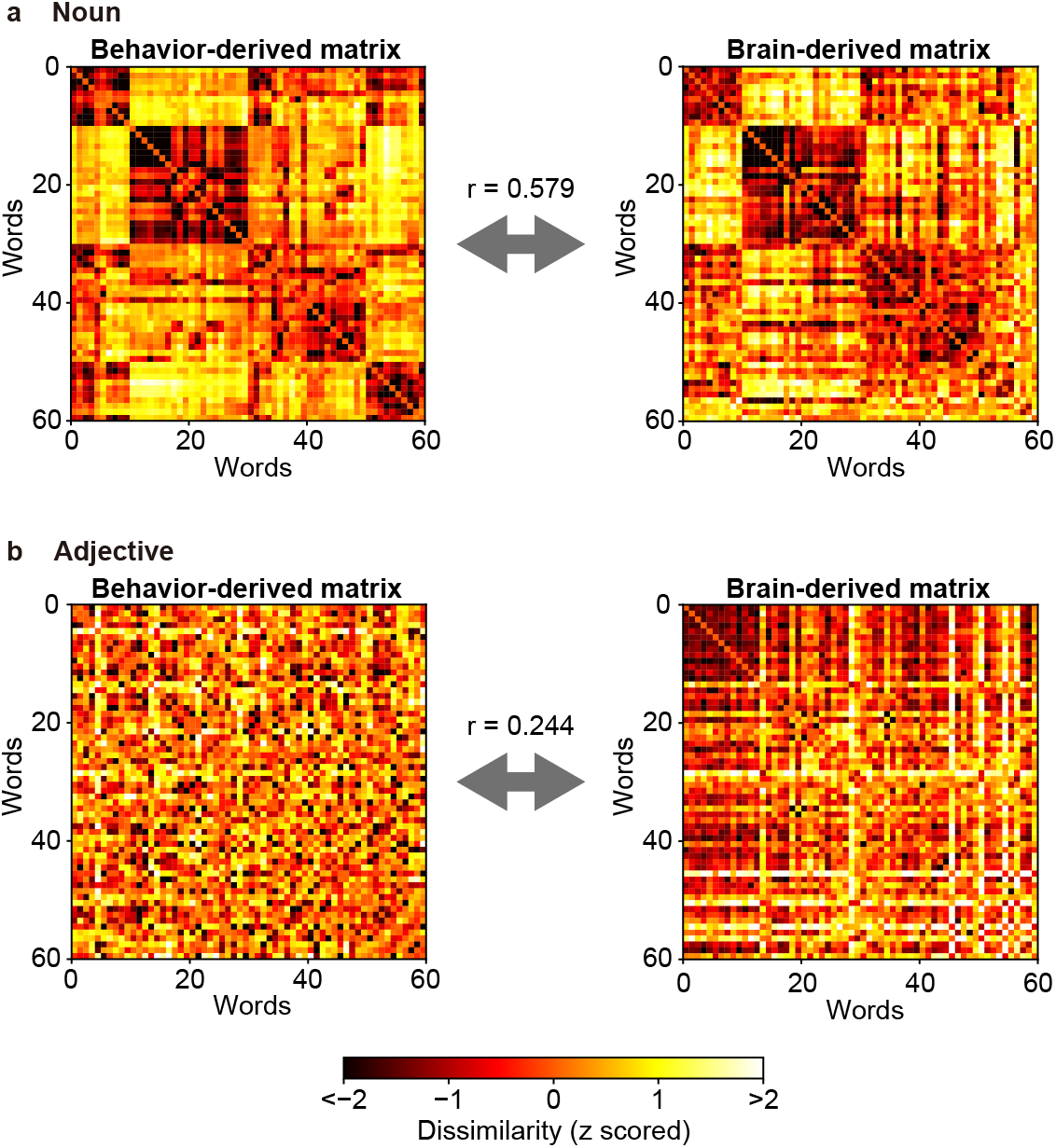
Correlations between brain- and behavior-derived dissimilarity matrices from the same participant population. We constructed word dissimilarity matrices separately for nouns (**a**) and adjective (**b**) using behavioral and voxelwise-model data obtained from the same population of 6 participants (word-vector dimension = 1000; movie set 1). The same conventions were used as in Fig 5. There were significant correlations between behavior- and brain-derived word dissimilarity matrices for both nouns and adjectives (p < 0.00001).

**S5 Fig.**
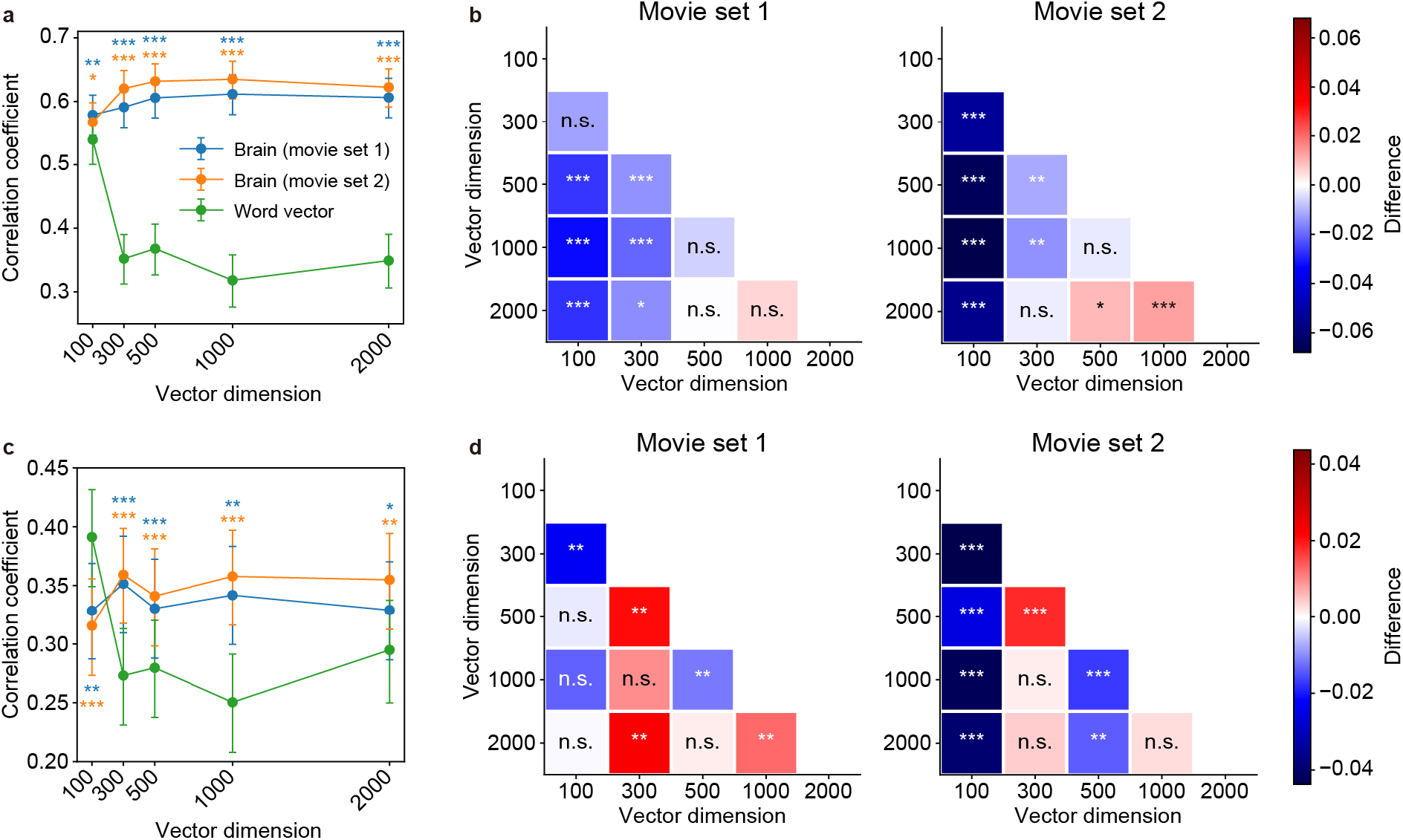
Correlations between dissimilarity matrices for all vector dimensions. **a**) Brain–behavior and word vector–behavior correlations of noun dissimilarity matrices for different vector dimensions. Error bars indicate 95% confidence interval (CI) estimated by bootstrapping. Marks above or below each plot indicate the statistical significance of the difference between brain–behavior and word vector–behavior correlation coefficients for each movie set (bootstrapping test, ***p < 0.001, **p < 0.01, *p < 0.05, FDR corrected) **b**) Difference between brain–behavior correlation coefficients of different vector dimensions for nouns. The color of each cell represents the coefficient difference of the dimension on the x-axis minus the dimension on the y-axis (red, positive values; blue, negative values). The mark in each cell indicates the statistical significance of the difference (bootstrapping test, ***p < 0.0001, **p < 0.01, *p < 0.05, n.s., p > 0.05, FDR corrected). (**c** and **d**) The same analyses were performed as in **a** and **b** using adjective dissimilarity matrices.

**S6 Fig.**
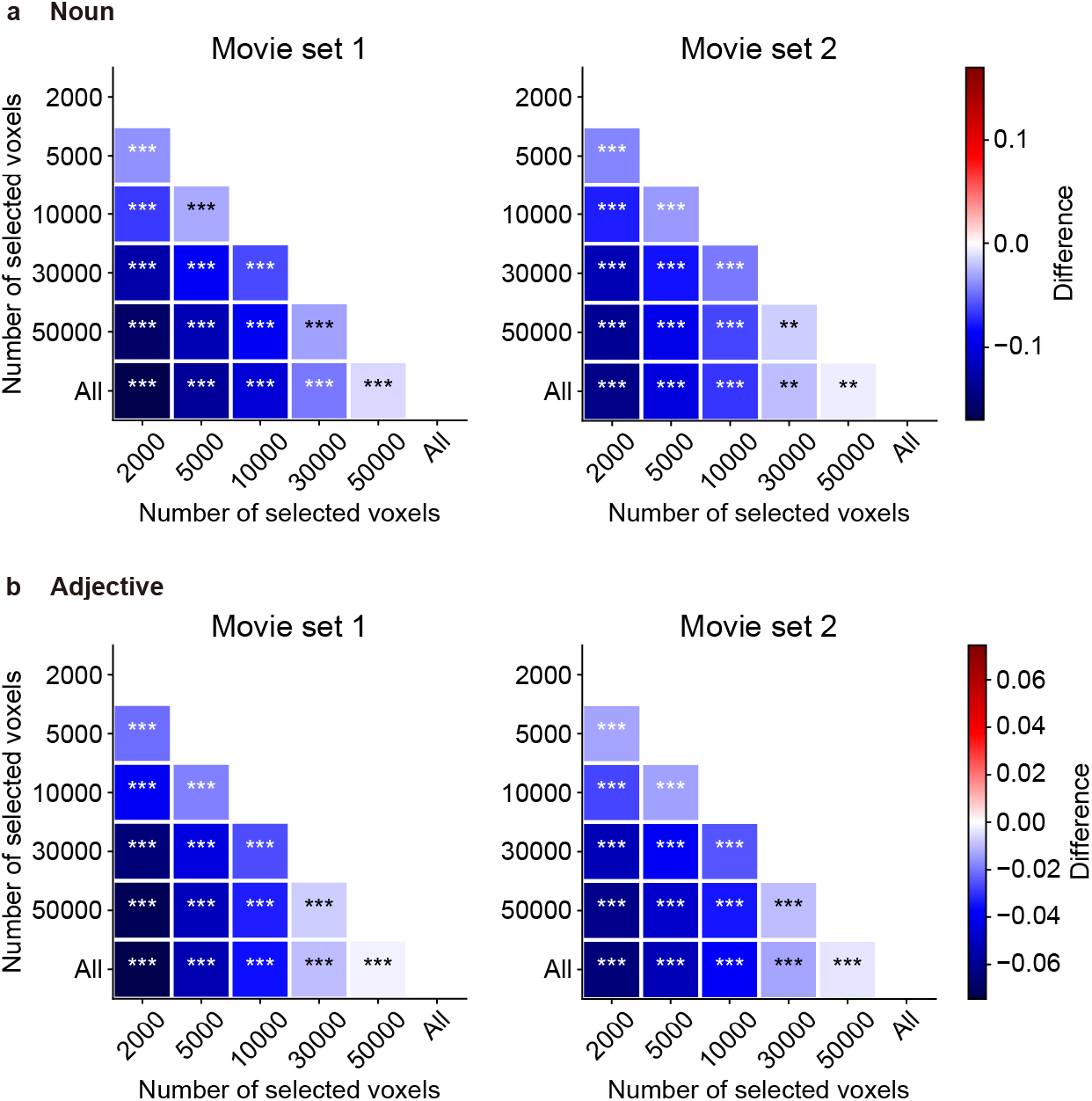
Difference of brain–behavior correlation coefficients between each pair of different voxel selections. The color of each cell represents the difference between each pair of the number of selected voxels; the difference is the brain–behavior correlation coefficient for the dimension on the x-axis minus the coefficient for the dimension on the y-axis (red, positive values; blue, negative values). The mark in each cell indicates the statistical significance of the difference (bootstrapping test, ***p < 0.0001, FDR corrected).

**S1 Table.**
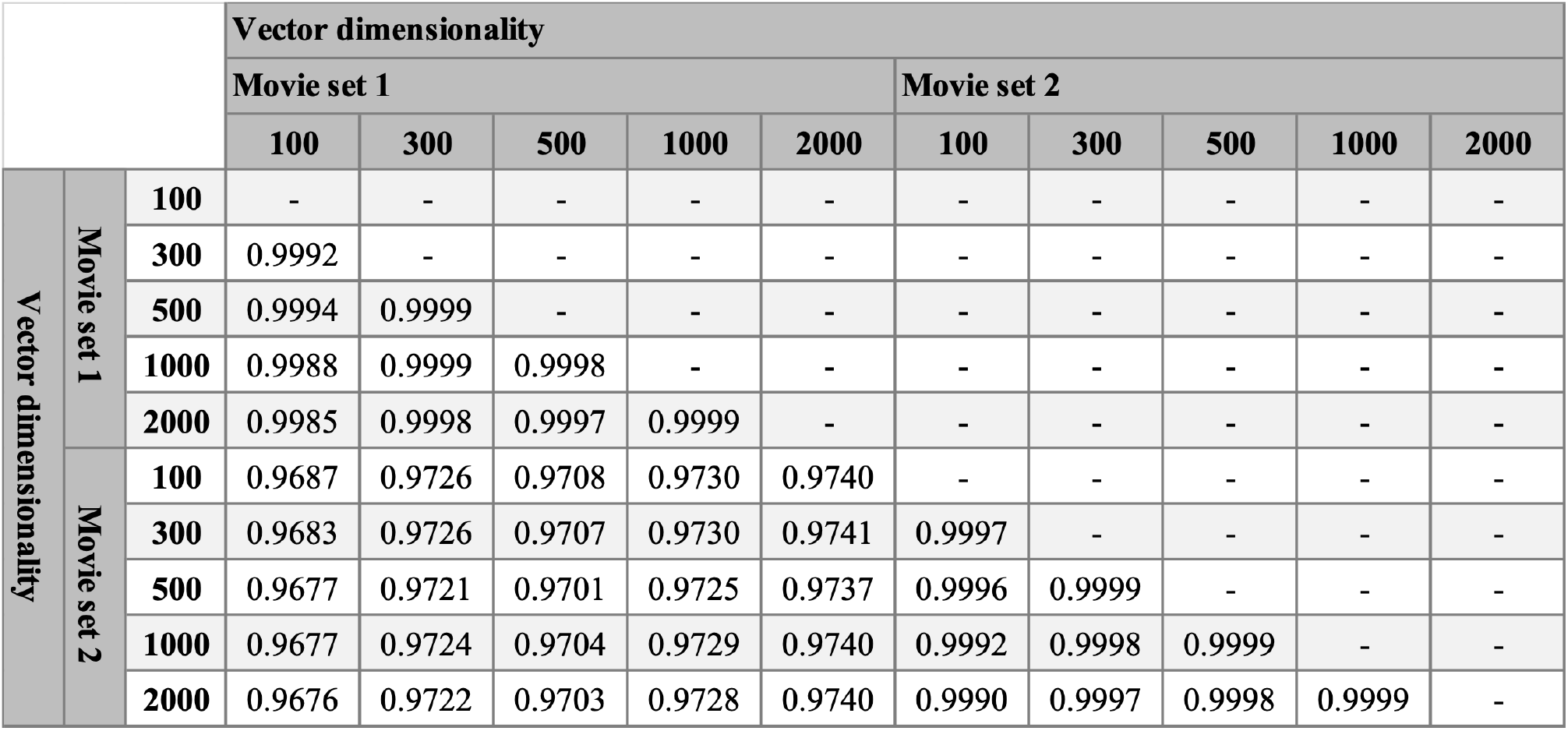
Consistency of inter-regional patterns of prediction accuracy across vector dimensionality and across movie sets. We calculated the mean prediction accuracy within each cortical region (Fig 4**a**–**b** and S2 Fig) and compared the inter-regional patterns of prediction accuracy between arbitrary pairs of vector dimensionality and movie sets. The value in each cell denotes the Pearson’s correlation coefficient of the inter-regional patterns between each pair.

**S2 Table.**
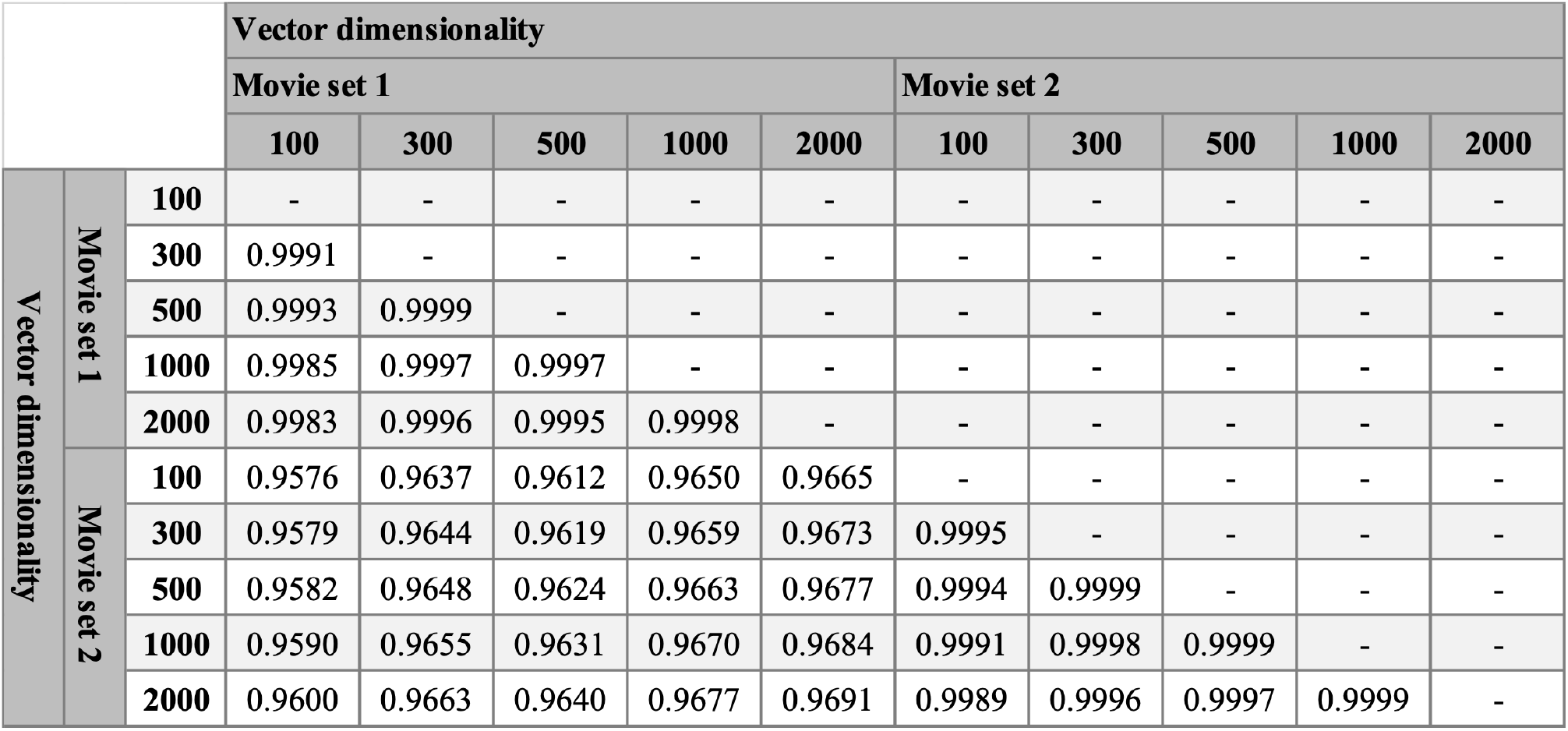
Consistency of inter-regional patterns of significant-voxel fraction across vector dimensionality and across movie sets. As in the case of the mean prediction accuracy (S1 Table), the inter-regional patterns of significant-voxel fraction (Fig 4**c**–**d** and S3 Fig) were compared between arbitrary pairs of vector dimensionality and movie sets. The value in each cell denotes the Pearson’s correlation coefficient of the inter-regional patterns between each pair.

**S3 Table.** Number of stimulus movie clips in individual product/service categories.

| **Categories** | **Movie set 1** | **Movie set 2** |
| --- | --- | --- |
| Electronic & Precision | 9 | 4 |
| Audiovisual | 1 | 5 |
| Appliance | 5 | 16 |
| Car | 33 | 31 |
| Food & Confectionery | 69 | 7 |
| Beverage & Alcoholic drink | 38 | 20 |
| Medical & Health | 21 | 35 |
| Cosmetics | 6 | 49 |
| Sundries & Home equipment | 45 | 10 |
| Garment/apparel | 11 | 9 |
| Entertainment | 43 | 42 |
| Media & Education | 16 | 41 |
| Distribution & Retailer | 20 | 12 |
| Communication & Service | 51 | 35 |
| House & Construction | 12 | 9 |
| Finance | 20 | 9 |
| Enterprise, Public service, & Others | 20 | 34 |

